# Identification of Pathogenic Structural Variants in Rare Disease Patients through Genome Sequencing

**DOI:** 10.1101/627661

**Authors:** James M. Holt, Camille L. Birch, Donna M. Brown, Manavalan Gajapathy, Nadiya Sosonkina, Brandon Wilk, Melissa A. Wilk, Rebecca C. Spillmann, Nicholas Stong, Hane Lee, Alden Y. Huang, Devon Bonner, Jennefer N. Kohler, Ellen F. Macnamara, Undiagnosed Diseases Network, Stanley F. Nelson, Vandana Shashi, Elizabeth A. Worthey

## Abstract

**Purpose:** Clinical whole genome sequencing is becoming more common for determining the molecular diagnosis of rare disease. However, standard clinical practice often focuses on small variants such as single nucleotide variants and small insertions/deletions. This leaves a wide range of larger “structural variants” that are not commonly analyzed in patients.

**Methods:** We developed a pipeline for processing structural variants for patients who received whole genome sequencing through the Undiagnosed Diseases Network (UDN). This pipeline called structural variants, stored them in an internal database, and filtered the variants based on internal frequencies and external annotations. The remaining variants were manually inspected and then interesting findings were reported as research variants to clinical sites in the UDN.

**Results:** Of 477 analyzed UDN cases, 286 cases (≈ 60%) received at least one structural variant as a research finding. The variants in 16 cases (≈ 4%) are considered “Certain” or “Highly likely” molecularly diagnosed and another 4 cases are currently in review. Of those 20 cases, at least 13 were identified originally through our pipeline with one finding leading to identification of a new disease. As part of this paper, we have also released the collection of variant calls identified in our cohort along with heterozygous and homozygous call counts. This data is available at https://github.com/HudsonAlpha/UDN_SV_export.

**Conclusion:** Structural variants are key genetic features that should be analyzed during routine clinical genomic analysis. For our UDN patients, structural variants helped solve ≈ 4% of the total number of cases (≈ 13% of all genome sequencing solves), a success rate we expect to improve with better tools and greater understanding of the human genome.

## Introduction

An individual rare disease affects fewer than 1 in 2000 people in the general population, but rare diseases are collectively common with an estimated 10% of the population having one or more rare diseases [1]. Many rare diseases are known to have underlying genetic causes and others, especially those affecting young children, are expected to have a genetic cause. In order to identify these genetic causes, genomic sequencing is becoming a common practice for the initial identification of candidate molecular causes [2, 3, 4].

The Undiagnosed Diseases Network (UDN) is a multi-site collaborative effort that uses genomic sequencing as one of the core resources for identifying molecular diagnoses of rare diseases [2]. Most patients who are accepted to the UDN are candidates for exome or genome sequencing, meaning they are believed to have a disorder caused or influenced by their genome. These patients are very diverse from a phenotype perspective, and the majority of them have been experiencing those phenotypes since birth or early childhood. Additionally, most patients have been through a wide range of prior genetic testing such as karyotyping, targeted gene testing, gene panels, and/or prior exome or genome sequencing that did not yield a molecular diagnosis.

To search for a molecular diagnosis, most accepted UDN participants are sequenced using exome sequencing or genome sequencing. The sequencing data is run through standard clinical pipelines at the sequencing cores that generally consist of alignment, variant calling, variant annotation, variant filtering, variant analysis, and reporting [5]. These standard clinical pipelines are targeted toward small variants such as single nucleotide variants and small (*≤*50bp) insertions or deletions. Unfortunately, variants that are too large for the variant callers (usually *>*50bp) are missed in this process.

Some variants, such as large deletions or duplications (*>*100kbp), are sometimes detected prior to genomic sequencing using targeted approaches or arrays. However, this still leaves a relatively large gap of unaccounted for variants ranging from 50bp to 100kbp. For this paper, we will consider “structural variants” to be deletions, duplications, insertions, translocations, and inversions that are larger than 50bp in size [6]. In recent years, advances in detecting structural variants from sequencing data has lead to an abundance of tools that are publicly available (e.g. [7, 8, 9, 10, 11, 12]) that can detect these structural variants and output them in a standard file format for downstream analyses.

Structural variant callers have already been used to help identify molecular diagnoses in patients with rare diseases. One group reported clinically significant copy-number variation in 15% of 79 trial rare, undiagnosed disease cases [13]. Another group searched for complex structural variation (defined as variation with three or more breakpoints/junctions), reporting three pathogenic complex structural variants out of 1324 patients with rare, undiagnosed disease [14]. Both groups relied on clinical orthogonal methods such as microarrays or Sanger sequencing to confirm the variants.

In this work, we developed a pipeline for analyzing structural variant calls in rare disease cases and then applied the pipeline to patients who received genome sequencing through the UDN. Our pipeline relies on the standard clinical pipeline for performing the sequencing and alignment steps. We then ran a structural variant caller with high sensitivity (Manta [10]) and filtered the resulting structural variant calls to a small set of rare calls that were then manually inspected for clinical relevance. Out of 477 UDN cases, we report that 20 cases (4.19%) were classified as “Certain” molecular diagnosis, “Highly likely” molecular diagnosis, or are in review as a result of this structural variant analysis. Finally, we generated a VCF file containing all observed structural variants in our cohort along with the number of times each variant was found.

## Material and Methods

### Genome Sequencing

For the Undiagnosed Diseases Network (UDN), we extracted DNA from whole blood samples and sequenced it using standard operating protocols that are validated for use as a Laboratory-Developed Test (LDT) in a CAP/CLIA lab [2]. A few samples were extracted from fibroblast samples instead of or in addition to blood samples when the clinical site deemed it appropriate. Approximately 480 million paired-end reads of 150bp in length were generated for each sample. When available and at the discretion of the clinical sites, family members were also sequenced in order to inform downstream analyses. After sequencing, all base calling and read filtering were performed with current Illumina software at the time of sequencing. We followed the GATK best practices [15] to align to the human reference genome (GRCh37) with BWA-mem [16]. The aligned sequences were then processed via GATK for base quality score recalibration, indel realignment, and duplicate removal [15].

### Structural Variant Calling

The main output of the previous step is one output BAM file per sample that contains all read mappings generated from alignment to the reference genome. We then ran a structural variant caller on that BAM file. While any structural variant caller could be used in this pipeline, we intentionally selected a caller with high sensitivity (i.e. a high rate of finding true structural variants). While precision is important as well, we instead relied on downstream filtering to remove common false positives from the call set.

For testing the structural variant callers, we downloaded the Genome in a Bottle high-confidence deletion dataset to use as our truth set against sample NA12878 [17]. At the time of initial pipeline development, *Manta* v1.0.1 [10] was found to have the highest sensitivity (91%) amongst the structural variant software we internally tested. Of note, we found Manta had a relatively low precision (52%), indicating that there are a relatively high number of false positives in its output. In practice, we found these false positives tend to be repeated across unrelated samples and are easy to remove due to their relatively high call rate. For a comprehensive evaluation of current structural variant callers, we recommend the analyses from *Parliament2* [12].

### Structural Variant Filtering

For a single sample, we found that Manta generally generates 10-14 thousand variant calls: 2-3 thousand break-end/translocation calls, 5-7 thousand deletions, *<*1 thousand duplications, 2-3 thousand insertions, and *<*1 thousand inversions. While there are several orders of magnitude fewer structural variant calls than small variant calls, manual inspection of each one is still impractical. To address this problem, we stored all variants in a database and developed a standard filter intended to identify candidate variants for any rare disease patient coming through the pipeline.

We designed this filter to mirror a clinical filter used for small variants, focusing on these expectations for a causal variant for a rare disease: 1) low allele frequency in the population, 2) impacting/overlapping an annotated gene, and 3) high quality variant call.

First, we expected deleterious variants to be rare in the general population. For a given proband, we filtered out variants with an allele frequency greater than 0.5% in our internal database at the time of analysis (note: initially the database contained approximately 500 samples, so the database frequencies changed over time as more samples arrived). In addition to filtering out true common variants, this first filter had the added benefit of filtering out common false positives such as those generated by upstream technical artifacts or the structural variant caller itself. Note that for the purpose of calculating internal allele frequency, we treated structural variants with imprecise breakpoints as identical if their breakpoint confidence intervals overlapped at all.

Second, we filtered out variants that do not overlap annotated genes from RefSeq [18]. The main reason for this filter is interpretation of the variant. Variants that do not directly impact a gene are currently difficult to clinically interpret, so they were removed from consideration during filtering.

Finally, we filtered out variant calls with low supporting evidence as described by the split read (SR) and paired read (PR) tags from Manta. For all variants, we required there to be at least eight total reads spanning the locus and for at least 15% of split reads to support the variant. The only exception to this requirement was for imprecise structural variant calls, for which Manta did not report any split reads. For larger events (breakends, deletions longer than 200bp, and duplications longer than 200 bp), we additionally required paired read support using the same total depth count (8) and variant support percentage (15%) criteria. After the above filtering, there were typically 20-50 variant calls remaining for a sample that were ready for variant analysis.

### Variant Analysis

For deletion, duplication, and inversion calls that were smaller than 1Mbp, we used a tool called *samplot* [19] to generate visualizations of the structural variant call for the proband and any available relatives in a case. These visualizations allowed analysts to see coverage changes and discordant read pairs for the proband and relatives at the site of the structural variant call of interest. If the coverage and discordant read pairs support the variant call, it passed manual inspection. In our experience, samplot tended to fail for variants larger than 1Mbp, so in-house visualizations were used for these larger structural variant calls.

For duplication and deletion calls smaller than 5Mbp, we generated basic coverage plots from the BAM files and checked for changes in coverage corresponding to the structural variant in question. Deletions and duplications larger than 5Mbp were trivially inspected using whole chromosome coverage plots. In our experience, the largest calls (*>*1Mbp) were almost always false positives. Those that were true positives were usually already noted by the clinical site through a prior test. Any variant calls that failed manual inspection were treated as false positives and removed from further consideration.

Variants that passed manual inspection were then analyzed for phenotypic relevance to the patient, similar to how a small variant would be considered in the standard clinical pipeline. This primarily consisted of reviewing genes impacted by the structural variant and then cross-checking those genes for known diseases in publicly available databases such as OMIM [20] or the Human Phenotype Ontology [21]. If a variant was deemed to impact a gene with a matching disease phenotype, it was sent back to the UDN clinical site as a research variant. Additionally, variants that were sufficiently “large” (*>*100kbp for deletions or *>*300kbp for duplications/inversions) were also returned provided that they passed the filtering and manual inspection process described above.

## Results

### Case Level Summary

At the time of writing, we have run Manta on 1641 UDN individuals (probands and relatives) who received genome sequencing, and we have analyzed structural variants for 477 UDN cases. Of those cases, 286 (59.96%) had at least one structural variant that was ultimately returned to the corresponding clinical site as a research variant. Table 1 shows a summary of how many variants were returned per case.

**Table 1:** Case Return Summary. This table shows the summary of how many structural variants were returned per case. Of note, ≈ 60% of cases had at least one structural variant returned as a research variant.

| Number of Returned Variants ($X$) | Number of Cases with $X$ Returned Variants |
| --- | --- |
| 1 | 180 (37.74%) |
| 2 | 79 (16.56%) |
| 3 | 17 (3.56%) |
| 4 | 7 (1.47%) |
| 5 | 3 (0.63%) |

We also categorized the reason a structural variant was returned into one or more of six categories: “phenotype” means the variant impacts a gene that has phenotypic relevance to the patient, “size” means the event was “large” (*>*100kbp for deletions or *>*300kbp for duplications/inversions), “*de novo*” means both parents were also sequenced and the variant was not called or detected via manual inspection in either, “homozygous” means the variant was called on both copies of the chromosome, “compound heterozygous” means the variant was found to be in *trans* with another variant (structural or small) impacting the same gene, and “hemizygous” means the patient only has one copy of the chromosome and that copy has the variant (i.e. variants on chromosome X for a male). Table 2 shows a breakdown of the returned structural variants by type and reason for return.

**Table 2:**
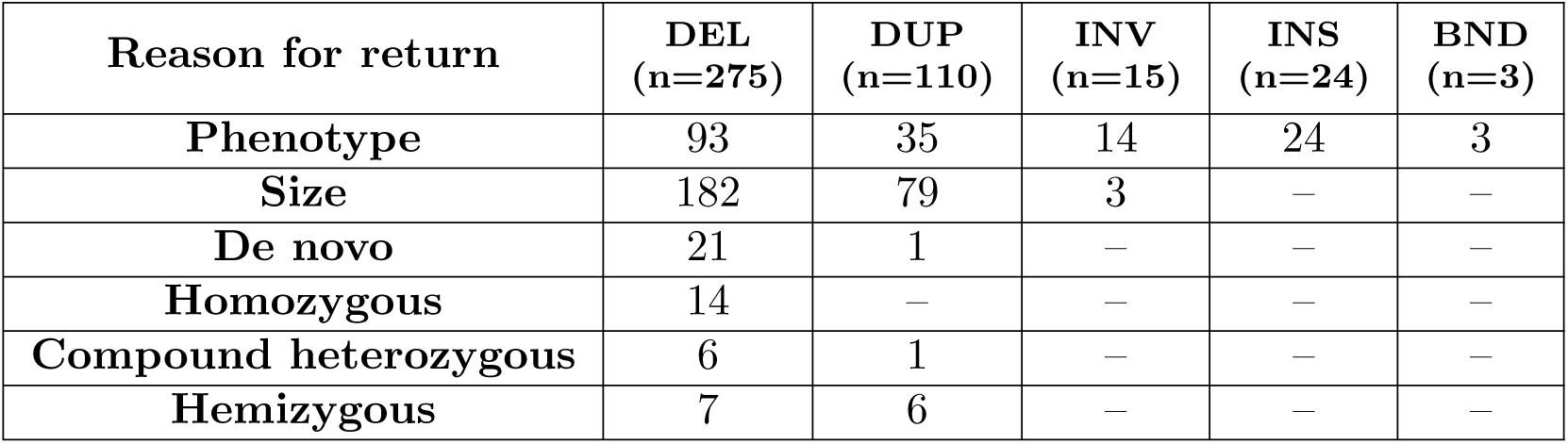
Returned Structural Variants Summary. This table shows the summary of structural variants that were returned to clinical sites in the UDN. On the top row is the type of variant that was returned and the total number of times that type was returned for deletions (DEL), duplications (DUP), inversions (INV), insertions (INS), and break-end pairs (BND). On the left column is the reason that a particular variant was returned. Note that an individual variant may have more than one reason for being returned.

| Reason for return | DEL<br>(n=275) | DUP<br>(n=110) | INV<br>(n=15) | INS<br>(n=24) | BND<br>(n=3) |
| --- | --- | --- | --- | --- | --- |
| Phenotype | 93 | 35 | 14 | 24 | 3 |
| Size | 182 | 79 | 3 | – | – |
| De novo | 21 | 1 | – | – | – |
| Homozygous | 14 | – | – | – | – |
| Compound heterozygous | 6 | 1 | – | – | – |
| Hemizygous | 7 | 6 | – | – | – |

The most returned variant type was deletions (n=275) followed by duplications (n=110). There are several likely explanations for why deletions were reported more than any other category. First, in our samples, deletions are the structural variant type that tends to get called the most in Manta, so there is likely some ascertainment bias from the variant caller. Second, a large portion of deletions were returned simply because they were “large”, rare events with no other indication of deleteriousness. Lastly, interpretation of deletions (especially those that disrupt coding regions) is usually based on haploinsufficiency, and may be more clinically relevant than the interpretations from other structural variant types. This likely led to an increase in the number of deletions returned simply because interpretation was comparatively uncomplicated. Size and phenotype were the two most common reasons for return and were proportionally similar for deletions and duplications (i.e. cases with “phenotype” reason ≈ half of cases with “size” reason). However, this same pattern didn’t hold for the other reasons for return. Of note, while 21 returned deletion events were determined to be *de novo*, only one returned duplication event was found to be *de novo*. Additionally, reasons for return that reflect an autosomal recessive disorder (“Homozygous” or “Compound heterozygous”) were found 20 times in deletions but only once in duplications.

### Molecular Diagnoses

Across all 477 analyzed cases, 20 cases (4.19%) received structural variant calls that we believe explain all or part of their patient’s phenotypes. Of those 20 cases, the structural variants in 16 cases are classified as “Certain” or “Highly likely” molecular diagnoses by the UDN, meaning they were confirmed by the clinical site and are currently considered to explain most or all of the patient’s phenotypes. Another four cases have a high overlap with the patient’s phenotypes, but are currently in review by the corresponding clinical sites. The variant(s), impacted genes, patient phenotypes, case type, clinical status, and confirmation method for all 20 cases are described in Table 3. At the time of this writing, the 16 diagnosed cases represent 13.44% of all diagnosed cases that received genome sequencing through the UDN.

**Table 3:**
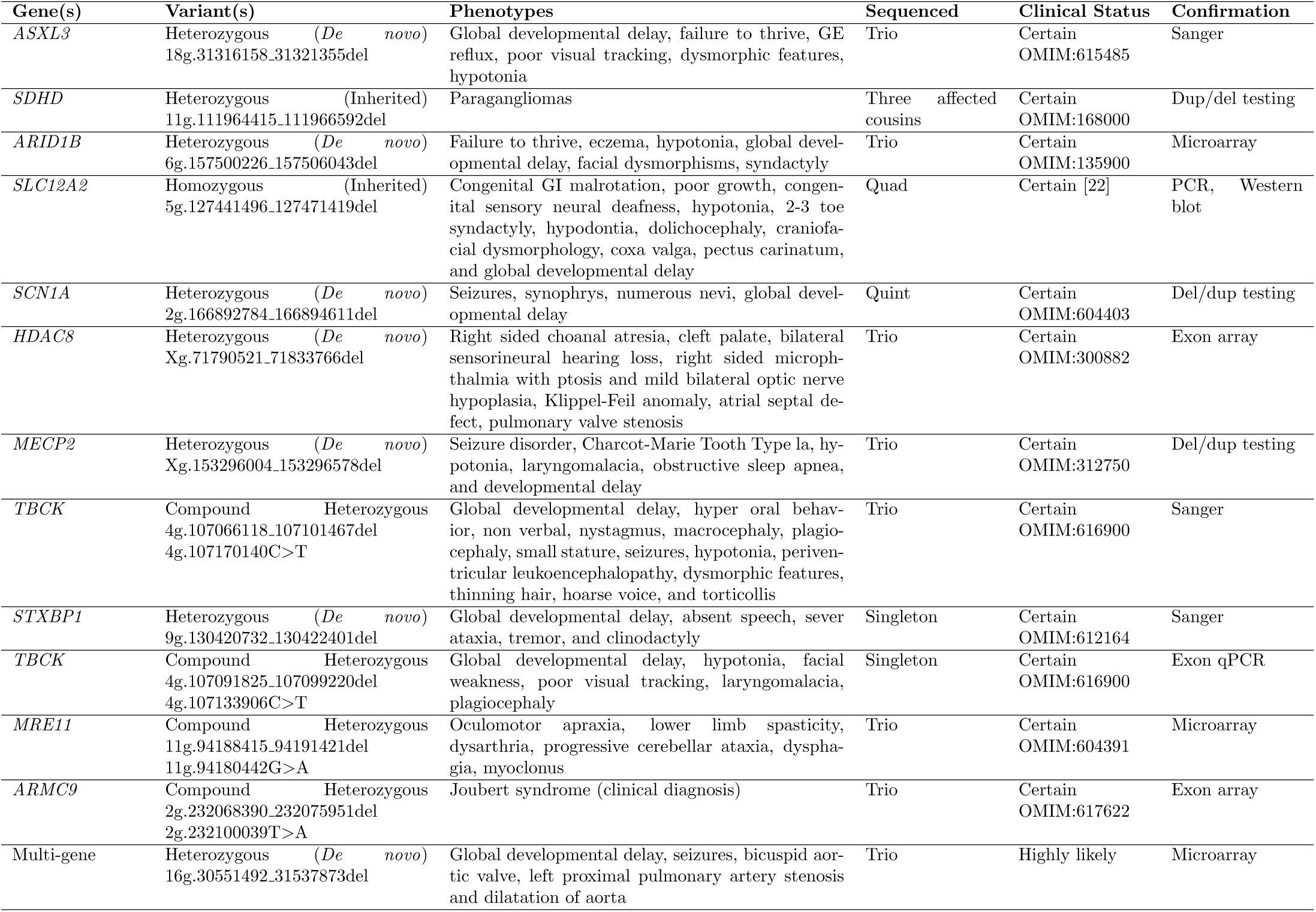
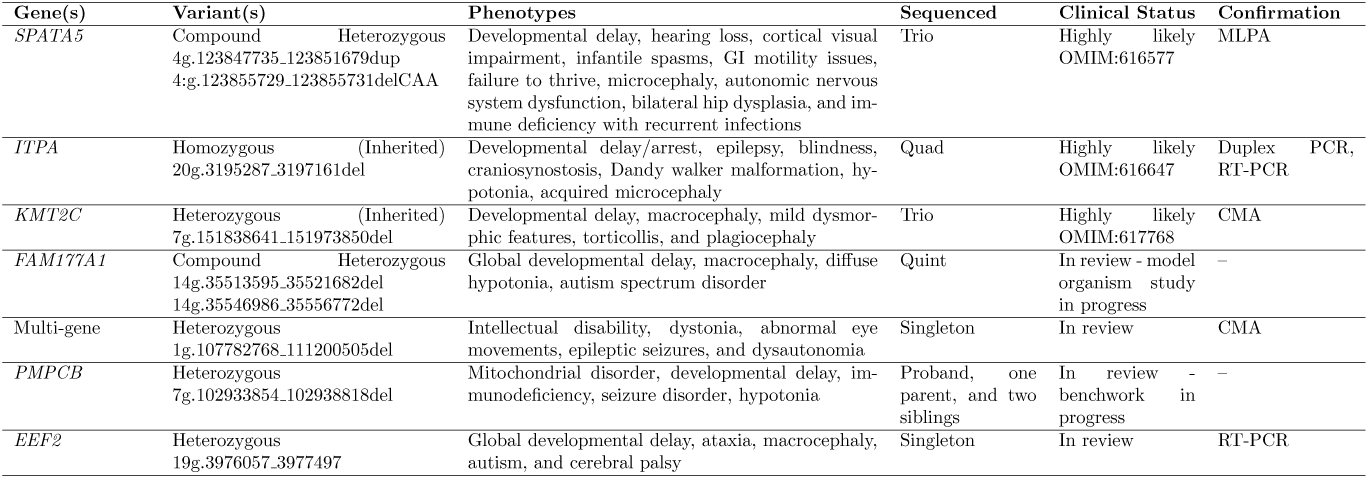
UDN Cases with Structural Variants. These are UDN cases with structural variants found using our pipeline that are either considered clinical solves or are still under review. For each case, we report the impacted genes, the zygosity of the variants, their genomic coordinates (GRCh37), major phenotypes, the sequenced family structure, clinical status, and CLIA confirmation status when available. When possible, we also note whether the variants are *de novo*, inherited, or compound heterozygous with another variant (all compound heterozygous variants were also inherited in our cases). For clinical status, we used official UDN statuses of “Certain” or “Highly likely”. Cases labeled as “In review” were considered clinically relevant by our team but have not been fully reviewed by the corresponding clinical site yet. As of this writing, twelve cases are certain solves, four cases are highly likely solves, and four cases are in various stages of review by the ordering clinical site.

Across all 20 cases, seven of the cases were reported as *de novo*, heterozygous variants in the proband. Six cases were reported as compound heterozygous, five of which are heterozygous with a reported small variant from the standard clinical pipeline and the sixth caused by two different inherited deletions impacting the same gene (case *FAM177A1* in Table 3). Two cases have homozygous, inherited variants. One of these cases with homozygous structural variants(case *ITPA* in Table 3) was caused by normal inheritance of the same structural variant from both parents. The other case (case *SLC12A2* in Table 3) was uniquely inherited from a single parent but was homozygous due to uniparental isodisomy in the patient. Additionally, the *SLC12A2* case was unique in that it led to the identification of a new disease described as Kilquist Syndrome [22].

To our knowledge, only the two large “Multi-gene” deletion variants were known by the clinical sites prior to running our pipeline. Additionally, five of the 20 cases would have likely received a partial diagnosis through the discovery of the small variant that was found in a compound heterozygous state. This leaves at least 13 cases that received a molecular diagnosis as a result of this structural variant pipeline. Of note, three of these cases (cases *SDHD, SCN1A*, and *MECP2* in Table 3) previously received negative results on one or more targeted tests on the impacted gene that was identified through our pipeline.

### Aggregate Variant Data

When we first developed our structural variant pipeline, one of the major resources we were lacking was a database of variant calls including allele counts and/or frequencies. In order to reduce this burden for other researchers, we have made available the structural variant call set that is stored in our database along with the number of times each variant was called across our 1641 UDN samples in a Variant Call Format (VCF) file. We stored variants in the format reported by Manta, including fields reflecting “imprecise” variants, confidence intervals for endpoint boundaries, and the variant length. For each variant, we reported the number of times that variant was called heterozygous and homozygous in our sample set. Note that due to imprecise variant calls and/or polymorphic sites, there are some variant calls that likely represent the same variant but were called differently in two or more samples. While our filtering process collapsed this imprecision into a single variant for the purpose of determining rarity, we intentionally left the calls separate to preserve the variant calls as Manta originally presented them. The end result is a VCF file containing 419657 deletions, 171401 duplications, 146873 inversions, 275885 insertions, and 654108 break-ends. This file is available at https://github.com/HudsonAlpha/UDN_SV_export.

## Discussion

In this work, we described a structural variant calling and filtering pipeline for rare diseases that was applied to patients in the Undiagnosed Diseases Network (UDN). In the 477 UDN cases analyzed, we found that 16 cases had variants that are considered molecular diagnoses (≈4% of all cases, ≈13% of all diagnoses) and another four cases are currently in review. Of these 20 cases, 13 were diagnosed as a direct result of our structural variant pipeline with one case representing a newly discovered disease. Additionally, three of these cases had previously received negative results on targeted gene tests for the gene that was impacted by the structural variant. Finally, approximately 60% of all cases had one or more structural variants that were returned as research variants that may warrant further investigation. Despite this relative success, we believe there is much room for improvement when it comes to identifying clinically relevant structural variants.

For our pipeline, we specifically selected a single structural variant caller with high sensitivity. While Manta was the best caller when this pipeline was originally conceived, methods that report better sensitivity and precision have since been created. For example, multi-caller systems such as Parliament2 [12] claim to increase accuracy by taking into account the output from multiple structural variant callers (including Manta). We expect that replacing Manta in our pipeline with a more accurate structural variant caller (or multi-caller system) will both increase the number of cases where relevant structural variants are identified and reduce the need to perform manual inspection of variants due to false positives. Additionally, each structural variant we identified needed to be CLIA confirmed through an orthogonal test before being returned to the patient. As the desire for clinical structural variants increases, we believe the need for CLIA certification of structural variant pipelines built on genome sequencing will become more important from both a time and cost perspective.

Another issue is that many of the annotations that are taken for granted in small variants are not readily available for structural variants. For example, when the pipeline was conceived, the main publicly available structural variant resources were dbVar [23] and DGV [24]. We found that the variants from either database rarely matched those that made it through our filtering process (anecdotally, *<*5% of filtered variants matched an entry in either database). Additionally, we had to rely on our own internal allele frequencies to determine the rarity of a variant call as there was no publicly available database of variant call frequencies. Fortunately, there are at least two large-scale resources for determining allele frequencies that are recently publicly available [25, 26], and we have made our raw structural variant call set available as a resource as well. Additionally, resources like ClinGen’s [27] dosage sensitivity map are now available to help in the interpretation of deletions and duplications. We expect that incorporating these new resources will improve our ability to identify clinically relevant structural variants, further improving the percentage of rare disease cases that can be molecularly diagnosed through sequencing and analysis of structural variants.

## Acknowledgements

This work was supported in part by the Intramural Research Program of the National Human Genome Research Institute and the NIH Common Fund through the Office of Strategic Coordination and Office of the NIH Director. Research reported in this manuscript was supported by the NIH Common Fund through the Office of Strategic Coordination and Office of the NIH Director under award numbers U01HG007530, U01HG007674, U01HG007703, U01HG007709, U01HG007672, U01HG007690, U01HG007708, U01HG007942, U01HG007943, U54NS093793, and U01TR001395. The content is solely the responsibility of the authors and does not necessarily represent the official views of the NIH.

## References

[1] Wright, C.F., FitzPatrick, D.R., Firth, H.V.: Paediatric genomics: diagnosing rare disease in children. Nature Reviews Genetics (2018)

[2] Ramoni, R.B., Mulvihill, J.J., Adams, D.R., Allard, P., Ashley, E.A., Bernstein, J.A., Gahl, W.A., Hamid, R., Loscalzo, J., McCray, A.T., et al.: The undiagnosed diseases network: accelerating discovery about health and disease. The American Journal of Human Genetics 100(2), 185–192 (2017)

[3] Bagnall, R.D., Ingles, J., Dinger, M.E., Cowley, M.J., Ross, S.B., Minoche, A.E., Lal, S., Turner, C., Colley, A., Rajagopalan, S., et al.: Whole genome sequencing improves outcomes of genetic testing in patients with hypertrophic cardiomyopathy. Journal of the American College of Cardiology 72(4), 419–429 (2018)

[4] Sweeney, N.M., Nahas, S.A., Chowdhury, S., Del Campo, M., Jones, M.C., Dimmock, D.P., Kingsmore, S.F., Investigators, R., et al.: The case for early use of rapid whole genome sequencing in management of critically ill infants: Late diagnosis of coffin-siris syndrome in an infant with left congenital diaphragmatic hernia, congenital heart disease and recurrent infections. Molecular Case Studies, 002469 (2018)

[5] Worthey, E.A.: Analysis and annotation of whole-genome or whole-exome sequencing derived variants for clinical diagnosis. Current protocols in human genetics 95(1), 9–24 (2017)

[6] Feuk, L., Marshall, C.R., Wintle, R.F., Scherer, S.W.: Structural variants: changing the landscape of chromosomes and design of disease studies. Human molecular genetics 15(suppl_1), 57–66 (2006)

[7] Abyzov, A., Urban, A.E., Snyder, M., Gerstein, M.: Cnvnator: an approach to discover, genotype, and characterize typical and atypical cnvs from family and population genome sequencing. Genome research 21(6), 974–984 (2011)

[8] Rausch, T., Zichner, T., Schlattl, A., Stütz, A.M., Benes, V., Korbel, J.O.: Delly: structural variant discovery by integrated paired-end and split-read analysis. Bioinformatics 28(18), 333–339 (2012)

[9] Zhu, M., Need, A.C., Han, Y., Ge, D., Maia, J.M., Zhu, Q., Heinzen, E.L., Cirulli, E.T., Pelak, K., He, M., et al.: Using erds to infer copy-number variants in high-coverage genomes. The American Journal of Human Genetics 91(3), 408–421 (2012)

[10] Chen, X., Schulz-Trieglaff, O., Shaw, R., Barnes, B., Schlesinger, F., Källberg, M., Cox, A.J., Kruglyak, S., Saunders, C.T.: Manta: rapid detection of structural variants and indels for germline and cancer sequencing applications. Bioinformatics 32(8), 1220–1222 (2015)

[11] Xi, R., Lee, S., Xia, Y., Kim, T.-M., Park, P.J.: Copy number analysis of whole-genome data using bic-seq2 and its application to detection of cancer susceptibility variants. Nucleic acids research 44(13), 6274–6286 (2016)

[12] Zarate, S., Carroll, A., Krasheninina, O., Sedlazeck, F.J., Jun, G., Salerno, W., Boerwinkle, E., Gibbs, R.: Parliament2: Fast structural variant calling using optimized combinations of callers. BioRxiv, 424267 (2018)

[13] Gross, A.M., Ajay, S.S., Rajan, V., Brown, C., Bluske, K., Burns, N.J., Chawla, A., Coffey, A.J., Malhotra, A., Scocchia, A., et al.: Copy-number variants in clinical genome sequencing: deployment and interpretation for rare and undiagnosed disease. Genetics in Medicine, 1 (2018)

[14] Sanchis-Juan, A., Stephens, J., French, C.E., Gleadall, N., Mégy, K., Penkett, C., Shamardina, O., Stirrups, K., Delon, I., Dewhurst, E., et al.: Complex structural variants in mendelian disorders: identification and breakpoint resolution using short-and long-read genome sequencing. Genome medicine 10(1), 95 (2018)

[15] DePristo, M.A., Banks, E., Poplin, R., Garimella, K.V., Maguire, J.R., Hartl, C., Philippakis, A.A., Del Angel, G., Rivas, M.A., Hanna, M., et al.: A framework for variation discovery and genotyping using next-generation dna sequencing data. Nature genetics 43(5), 491 (2011)

[16] Li, H.: Aligning sequence reads, clone sequences and assembly contigs with bwa-mem. arXiv preprint arXiv:1303.3997 (2013)

[17] Parikh, H., Mohiyuddin, M., Lam, H.Y., Iyer, H., Chen, D., Pratt, M., Bartha, G., Spies, N., Losert, W., Zook, J.M., et al.: svclassify: a method to establish benchmark structural variant calls. BMC genomics 17(1), 64 (2016)

[18] O’Leary, N.A., Wright, M.W., Brister, J.R., Ciufo, S., Haddad, D., McVeigh, R., Rajput, B., Robbertse, B., Smith-White, B., Ako-Adjei, D., et al.: Reference sequence (refseq) database at ncbi: current status, taxonomic expansion, and functional annotation. Nucleic acids research 44(D1), 733–745 (2015)

[19] Belyeu, J.R., Nicholas, T.J., Pedersen, B.S., Sasani, T.A., Havrilla, J.M., Kravitz, S.N., Conway, M.E., Lohman, B.K., Quinlan, A.R., Layer, R.M.: Sv-plaudit: A cloud-based framework for manually curating thousands of structural variants. bioRxiv, 265058 (2018)

[20] Amberger, J.S., Bocchini, C.A., Schiettecatte, F., Scott, A.F., Hamosh, A.: Omim. org: Online mendelian inheritance in man (omim R), an online catalog of human genes and genetic disorders. Nucleic acids research 43(D1), 789–798 (2014)

[21] Köhler, S., Carmody, L., Vasilevsky, N., Jacobsen, J.O.B., Danis, D., Gourdine, J.-P., Gargano, M., Harris, N.L., Matentzoglu, N., McMurry, J.A., et al.: Expansion of the human phenotype ontology (hpo) knowledge base and resources. Nucleic acids research 47(D1), 1018–1027 (2018)

[22] Macnamara, E., Koehler, A., D’Souza, P., Estwick, T., Lee, P., Vezina, G., Fauni, H., Braddock, S., Torti, E., Holt, J., et al.: Kilquist syndrome: A novel syndromic hearing loss disorder caused by homozygous deletion of slc12a2. Human mutation (2019)

[23] Lappalainen, I., Lopez, J., Skipper, L., Hefferon, T., Spalding, J.D., Garner, J., Chen, C., Maguire, M., Corbett, M., Zhou, G., et al.: Dbvar and dgva: public archives for genomic structural variation. Nucleic acids research 41(D1), 936–941 (2012)

[24] MacDonald, J.R., Ziman, R., Yuen, R.K., Feuk, L., Scherer, S.W.: The database of genomic variants: a curated collection of structural variation in the human genome. Nucleic acids research 42(D1), 986–992 (2013)

[25] Abel, H.J., Larson, D.E., Chiang, C., Das, I., Kanchi, K.L., Layer, R.M., Neale, B.M., Salerno, W.J., Reeves, C., Buyske, S., et al.: Mapping and characterization of structural variation in 17,795 deeply sequenced human genomes. bioRxiv, 508515 (2018)

[26] Collins, R., Brand, H., Karczewski, K., Zhao, X., Alföldi, J., Khera, A., Francioli, L., Gauthier, L., Wang, H., Watts, N., et al.: An open resource of structural variation for medical and population genetics (2019)

[27] Rehm, H.L., Berg, J.S., Brooks, L.D., Bustamante, C.D., Evans, J.P., Landrum, M.J., Ledbetter, D.H., Maglott, D.R., Martin, C.L., Nussbaum, R.L., et al.: Clingen?the clinical genome resource. New England Journal of Medicine 372(23), 2235–2242 (2015)

